# MetFish: A Metabolomics Platform for Studying Microbial Communities in Chemically Extreme Environments

**DOI:** 10.1101/518647

**Authors:** Chengdong Xu, Sneha P. Couvillion, Ryan L. Sontag, Nancy G. Isern, Yukari Maezato, Stephen R. Lindemann, Taniya Roy Chowdhury, Rui Zhao, Beau R. Morton, Ronald J. Moore, Janet K. Jansson, Vanessa L. Bailey, Paula J. Mouser, Margaret F. Romine, James F. Fredrickson, Thomas O. Metz

**Affiliations:** Earth and Biological Sciences Directorate, Pacific Northwest National Laboratory, Richland, WA, USA; Department of Civil and Environmental Engineering, University of New Hampshire, Durham, NH, USA

## Abstract

Metabolites have essential roles in microbial communities, including as mediators of nutrient and energy exchange, cell-to-cell communication, and antibiosis. However, detecting and quantifying metabolites and other chemicals in samples having extremes in salt or mineral content using liquid chromatography-mass spectrometry (LC-MS)-based methods remains a significant challenge. Here we report a facile method based on *in situ* chemical derivatization followed by extraction for analysis of metabolites and other chemicals in hypersaline samples, enabling for the first time direct LC-MS-based exo-metabolomics analysis in sample matrices containing up to 2 molar total dissolved salts. The method, MetFish, is applicable to molecules containing amine, carboxylic acid, carbonyl, or hydroxyl functional groups, and can be integrated into either targeted or untargeted analysis pipelines. In targeted analyses, MetFish provided limits of quantification as low as 1 nM, broad linear dynamic ranges (up to 5-6 orders of magnitude) with excellent linearity, and low median inter-day reproducibility (e.g. 2.6%). MetFish was successfully applied in targeted and untargeted exo-metabolomics analyses of microbial consortia, quantifying amino acid dynamics in the exo-metabolome during community succession; *in situ* in a native prairie soil, whose exo-metabolome was isolated using a hypersaline extraction; and in input and produced fluids from a hydraulically fractured well, identifying dramatic changes in the exo-metabolome over time in the well.

---

Microbial communities are ubiquitous and colonize a wide range of habitats and organisms, often thriving even in extreme environments with physicochemical conditions unsuitable for most other life forms. There is increasing evidence that microbial communities are responsible for a wide range of processes critical to the health of the ecosystems they inhabit and impact it in ways for which we currently have limited knowledge. Thriving in complex or extreme environments requires specific adaptations; therefore, studying these organisms lends evolutionary insight into microbial stress responses.^1, 2^ The balance between cooperation and competition in harsh conditions contributes to the resistance and resilience of these communities,^3–7^ and elucidating the role of chemical exchange and communication among members will provide an improved understanding of the underlying molecular mechanisms that might be exploited, as well as in the identification of beneficial natural products.^8–14^ While metagenomics studies have been conducted to identify genes encoding novel biosynthetic pathways^15–17^, the measurement of primary and secondary metabolites in chemically extreme environments has been hampered by the complexities of the associated sample matrices.

Mass spectrometry is an indispensable analytical tool for identifying, quantifying and structurally characterizing chemical and biological molecules with high sensitivity and accuracy.^18–21^ As the central workhorse for proteomics and metabolomics, liquid chromatography coupled with mass spectrometry (LC-MS) has played a critical role in the development of omics technologies that have enabled high throughput, systems biology investigations of organisms.^22–24^ However, performing exo-metabolomics analyses in environmental samples can be challenging, due to the complexity of the associated sample matrices. A particular challenge is the presence of high (e.g. mM to M) concentrations of salts and minerals, which can compromise the extraction of metabolites from the samples, and suppress the ionization of metabolites during LC-MS analysis, resulting in diminished or skewed quantitative performance.^25–27^ Until now, samples consisting of or derived from such matrices have precluded the application of LC-MS-based measurements of metabolites and other small molecules.

To address this, we present MetFish, a method based on chemical tagging and extraction for comprehensive and quantitative measurement of metabolites and other small molecules in LC-MS-prohibitive matrices. Named for its ability to selectively fish metabolites of interest from sample matrices based upon common functional groups, MetFish is comprised of four simple and inexpensive chemical tags targeting amine, carboxyl, carbonyl, and hydroxyl functional groups and allows for sensitive quantification of low abundance metabolites in both targeted and untargeted approaches. The four functional groups targeted by MetFish represent over 89% and 83% of the metabolites contained in the *E. coli* Metabolome and Plantcyc databases, respectively.^28, 29^ The chemical tags can be either used in tandem for untargeted global analysis of the metabolome or individually to profile the sub-metabolome by targeting the molecules containing a specific functional group. MetFish uses low cost, commercially-available reagents that 1) could be used by researchers with diverse skill sets studying myriad sample types; 2) facilitate physical separation of metabolites from salt, mineral and other matrix components that interfere with quantitative LC-MS-based analysis; and 3) can be deployed *in situ* to minimize sample manipulation.

We demonstrate the utility and simplicity of MetFish in LC-MS-based exo-metabolomics analyses of three samples containing or derived from microbial communities from diverse ecosystems: a hypersaline aquatic microbial community, a prairie soil, and fluids injected into and produced from a hydraulically fractured well, each consisting of or derived from hypersaline (i.e. from 400 mM to 2 M) sample matrices. MetFish demonstrated excellent sensitivity, reproducibility, and linear dynamic range, and is a simple, rapid and effective approach for addressing the needs of the broader research community.

## Results

### Background and Overview of MetFish

In our search for an effective and simple approach to separate metabolites from interfering matrix constituents such as high concentrations of salts, we evaluated several commercially available solid phase extraction (SPE) chemistries to capture metabolites from a hypersaline matrix (e.g. 2 M total dissolved salts) but all were unsuccessful (**Supplemental Table S1**). We determined that separation methods based on molecular weight (e.g., dialysis or size exclusion) were not suitable, since the masses of low molecular weight metabolites (e.g. glycine: 75.07 g/mol) overlap with those of salt components (e.g sulphate: 96.06 g/mol), resulting in loss of metabolites in the lower mass range. Subsequently, we explored chemical tagging and capture techniques, including metabolite enrichment by tagging and proteolytic release (METPR) and a derivatization approach developed by Mattingly et al.^30,31^ Both approaches were time consuming and required significant solid/liquid phase chemical synthesis (e.g. up to 1 week for a single METPR probe for a researcher with basic organic synthesis skills) for preparing the capture or derivatization reagents. Moreover, these techniques were not amenable for the *in situ* capture of metabolites. Recognizing the need for a more efficient method that could be readily adopted by researchers from a broad range of disciplines, we adopted a suite of dansylated and related reagents coupled with downstream enrichment. The reagents were selected for their low cost, commercial availability and ease of use to increase accessibility of the method in the research community. Dansylation has been used for decades as a derivatization method for quantification of amino acids based on fluorescence detection.^32^ More recently, Li and colleagues have used dansylated and related reagents for targeted profiling of various sub-metabolomes using LC-MS.^33–36^ We postulated that the derivatization chemistries associated with these reagents would be successful when applied in hypersaline matrices, and that we could then efficiently extract derivatized molecules from the samples and away from interfering salts. For MetFish, we selected dansylchloride, dansylhydrazine, dansylcadaverine, and 4-(dimethylamino)benzoyl chloride to specifically tag metabolites containing amine, carbonyl, carboxyl, and hydroxyl functional groups, respectively (**Fig. 1a**). The one-step derivatization reactions require as little as 10 minutes to a maximum of 120 minutes to couple the target metabolite (the ‘fish’) and the chemical tag (the ‘hook’), thus increasing its hydrophobicity and facilitating its extraction with organic solvent (the ‘line’) and concomitant enrichment from interfering components of the sample matrix (**Fig. 1b**). The tagged and extracted metabolites are subsequently analyzed using reversed phase liquid chromatography (LC) coupled with MS.^33–36^ The reversed-phase LC includes inline solid phase extraction, which focuses the tagged metabolites prior to the analytical separation and separates them from any residual matrix components. Tandem MS (MS/MS) is used to fragment the tagged metabolites, resulting in fragment ions that are uniform for a given reagent and unique for a given metabolite,^35^ providing identification confidence and metabolite specificity, respectively.

**Figure 1.**
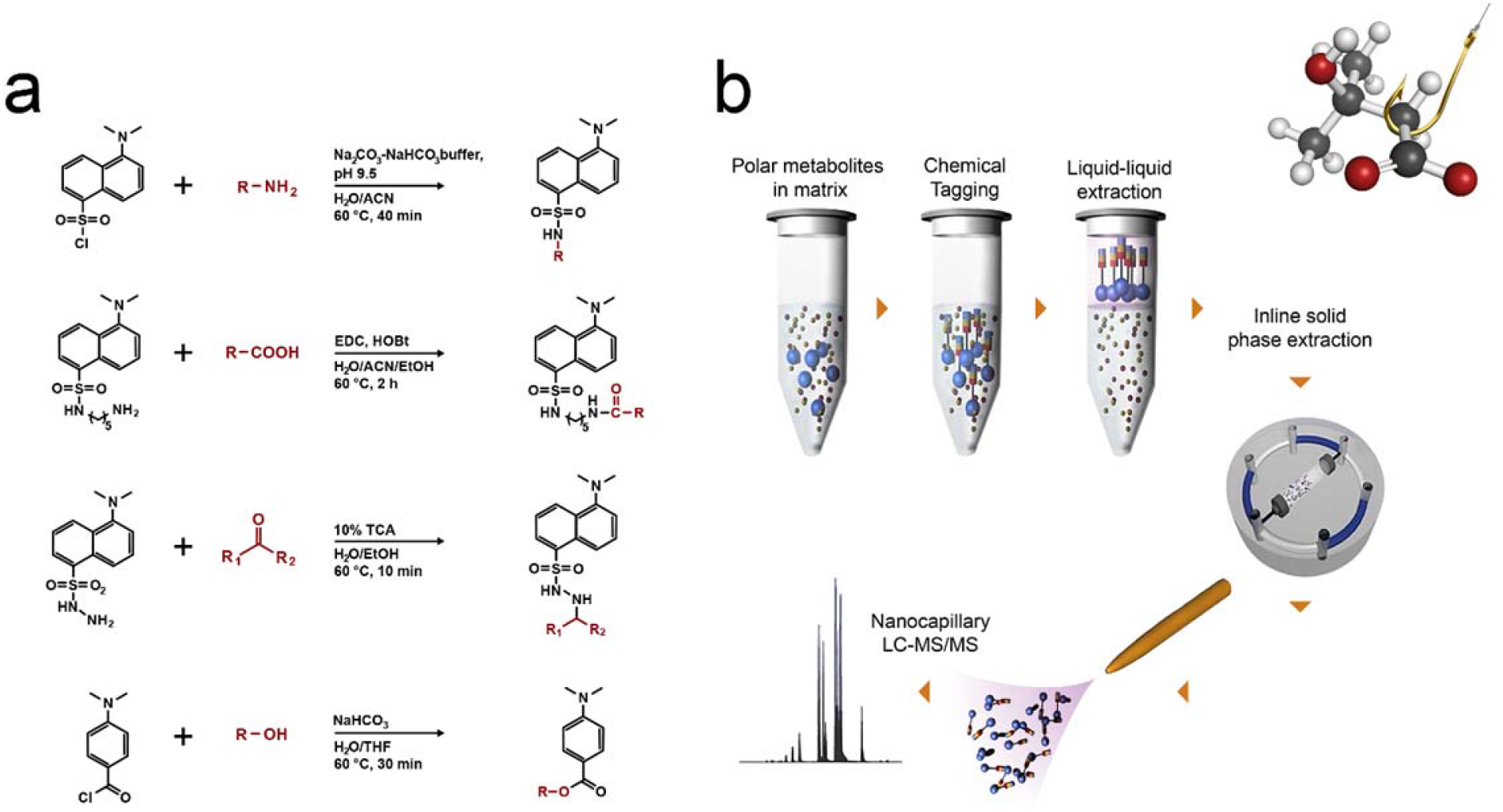
Overview of the MetFish method. **(a)** MetFish reagents and associated derivatization reactions **(b)** General workflow of the MetFish method

Exceptions to the latter are some isomeric metabolites, such as leucine and isoleucine, which do not produce unique fragment ions during collision-induced dissociation. To illustrate this, the fragmentation spectrum for dansylated glycine is shown in **Fig. 2a**. Fragment ions due only to the dansyl moiety are e.g. *m/z* 157, 170, and 252, whereas fragment ions due to dansyl-glycine are *m/z* 263 and 294. Some amount of the molecular ion *(m/z* 308) also appears in the MS/MS spectrum. All metabolites that have been tagged using the dansyl chloride reagent will generate the same fragment ions (e.g. *m/z* 157, 170, and 234, and 252), providing confidence in detection of an appropriately tagged amine-containing metabolite. In contrast, each dansylated metabolite will also generate fragment ions that are specific to the dansyl-metabolite complex and proportional in *m/z* to the mass of the tagged metabolite. The other MetFish reagents also produce uniform and specific fragment ions upon dissociation (**Supplemental Table S2**). These chemical characteristics enable MetFish reagents to be effective for both targeted and untargeted metabolomics applications. An added benefit is that differentially isotopically-labeled reagents can be used, allowing for the multi-plexing of labeled samples in untargeted metabolomics analysis, analogous to the iTRAQ and TMT peptide labeling approaches commonly used for multiplexing proteomics sample analyses using LC-MS/MS.^37^ Differences in abundances of “reporter ions” from MS/MS fragmentation of differentially labeled reagent-metabolite complexes would be used to provide accurate relative or absolute metabolite quantification. Alternatively, labeled metabolites could be incorporated as internal standards in targeted metabolite analysis.^33–36^ As shown in **Fig. 2a**, dansylated-^13^C and ^15^N-glycine produces fragment ions specific to the dansyl-glycine complex and with mass shifts proportional to the degree and type of isotope labeling.

**Figure 2.**
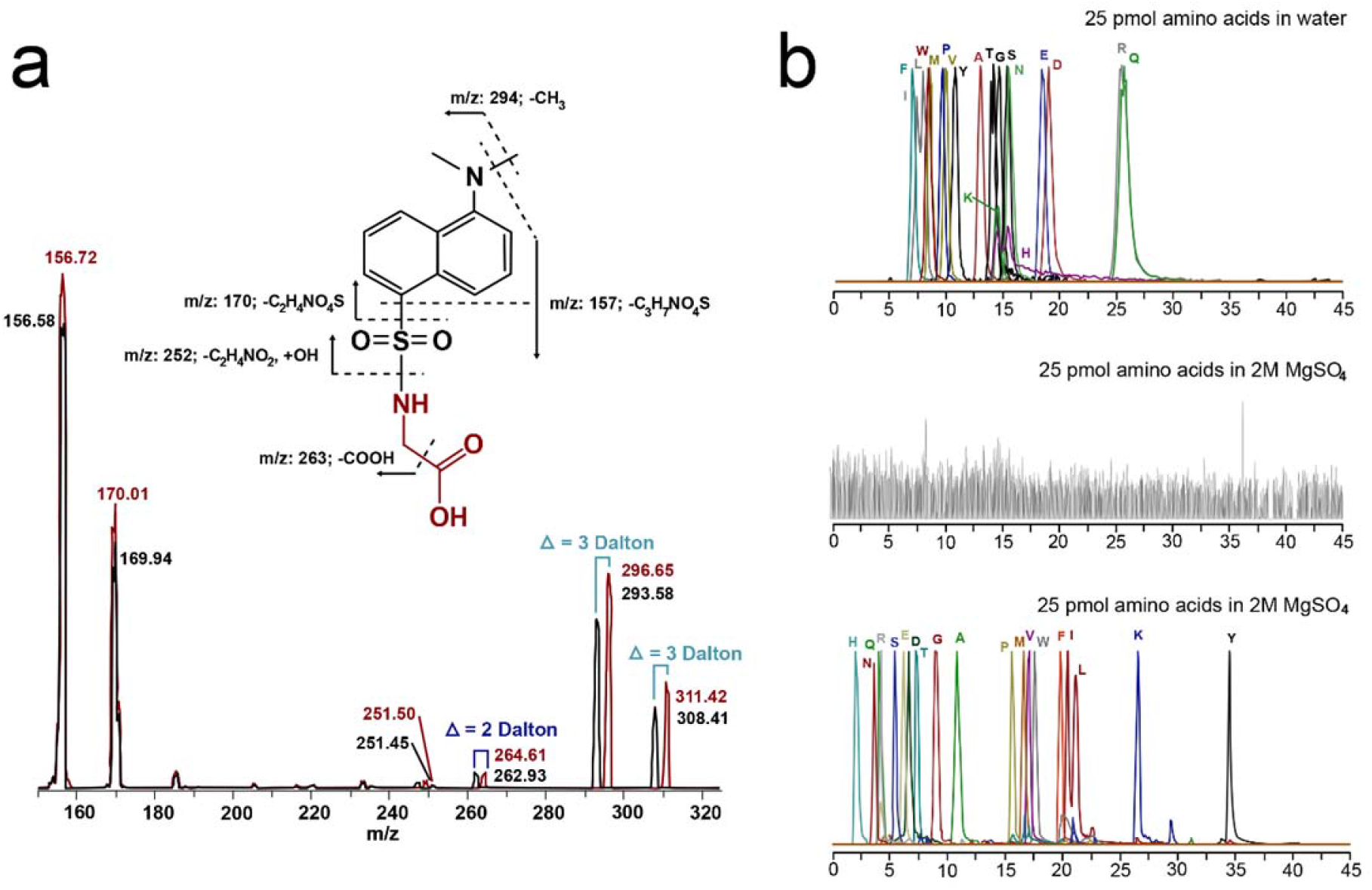
Validation of the MetFish method using amino acids. **(a)** Tandem mass spectra from analysis of a mixture of unlabeled (black spectrum) and ^13^C and ^15^N-labeled glycine (red spectrum), both derivatized with dansyl chloride. The *m/z* of each fragment peak is listed, and the mass shifts due to the isotopic labels are indicated. **(b)** Amino acids in neat solution analyzed by nanocapillary LC-MS/MS without chemical tagging (upper chromatogram); amino acids in 2 M MgSO_4_ analyzed by nanocapillary LC-MS/MS without chemical tagging (middle chromatogram); amino acids in 2 M MgSO_4_, derivatized using dansyl chloride, followed by extraction with organic solvent, and analyzed by nanocapillary LC-MS/MS with dansylation chemical tagging (lower chromatogram).

### Validation of MetFish

To assess the effectiveness of MetFish for targeted metabolite analysis in MS-prohibitive samples, we analyzed a mixture of 19 proteinogenic amino acids in water containing 2 M MgSO_4_, with and without the MetFish method and using LC-MS/MS with the mass spectrometer operating in selected reaction monitoring (SRM) mode. MgSO_4_ was chosen as it is a major salt component of Hot Lake, located in Oroville, WA, where a photoautotrophic microbial mat community resides and is available for study.^38, 39^ In typical MS-based metabolomics analyses, amino acids would be enriched from samples using extraction with organic solvents or a solid phase. As described above and shown in **Supplemental Table S1**, SPE is not effective for extracting small polar molecules from matrices containing high salt concentrations. Liquid / liquid extraction of amino acids from high-salt matrices either carries over sufficient salt in the extract to cause ionization suppression during analysis or does not effectively extract amino acids due to formation of amino acid-salt complexes that are insoluble in the organic solvent. As shown in **Fig. 2b**, top panel, analysis of a 25 pmol mixed amino acid standard dissolved in deionized water was straightforward using hydrophilic interaction liquid chromatography (HILIC)-MS/MS; however, no signal was observed above background for the same 25 pmol mixed amino acid standard dissolved in 2 M MgSO_4_ (**Fig. 2b**, middle panel). Applying the MetFish method using the amine tagging reagent resulted in quantitative measurement of all amino acids using reversed-phase LC-MS/MS (**Fig. 2b**, lower panel). Because of the increased hydrophobicity of the tagged amino acids, their SRM signals were also more intense (due to enhanced electrospray ionization^40^) and they were better resolved chromatographically using reversed-phase LC compared to their native forms measured using HILIC. In the MetFish analyses the unique fragment ion from each singly charged, tagged amino acid was used for quantification purposes, and a fragment ion common to all tagged amino acids (e.g. *m/z* 157, 170, or 252) provided confident identification.

To demonstrate the broad applicability of the MetFish approach for detecting metabolites containing other functional groups, we analyzed metabolites containing carbonyl, carboxyl, and hydroxyl functional groups. As with amino acids (**Figs. 2a and 3a**), the MetFish method enabled quantification of metabolites with carboxylic acids (**Fig. 3b**), carbonyl (**Fig. 3c**), and hydroxyl groups, including sugars (**Fig. 3d**) and alcohols (**Fig. 3e**), all in water containing 2 M MgSO4.

**Figure 3.**
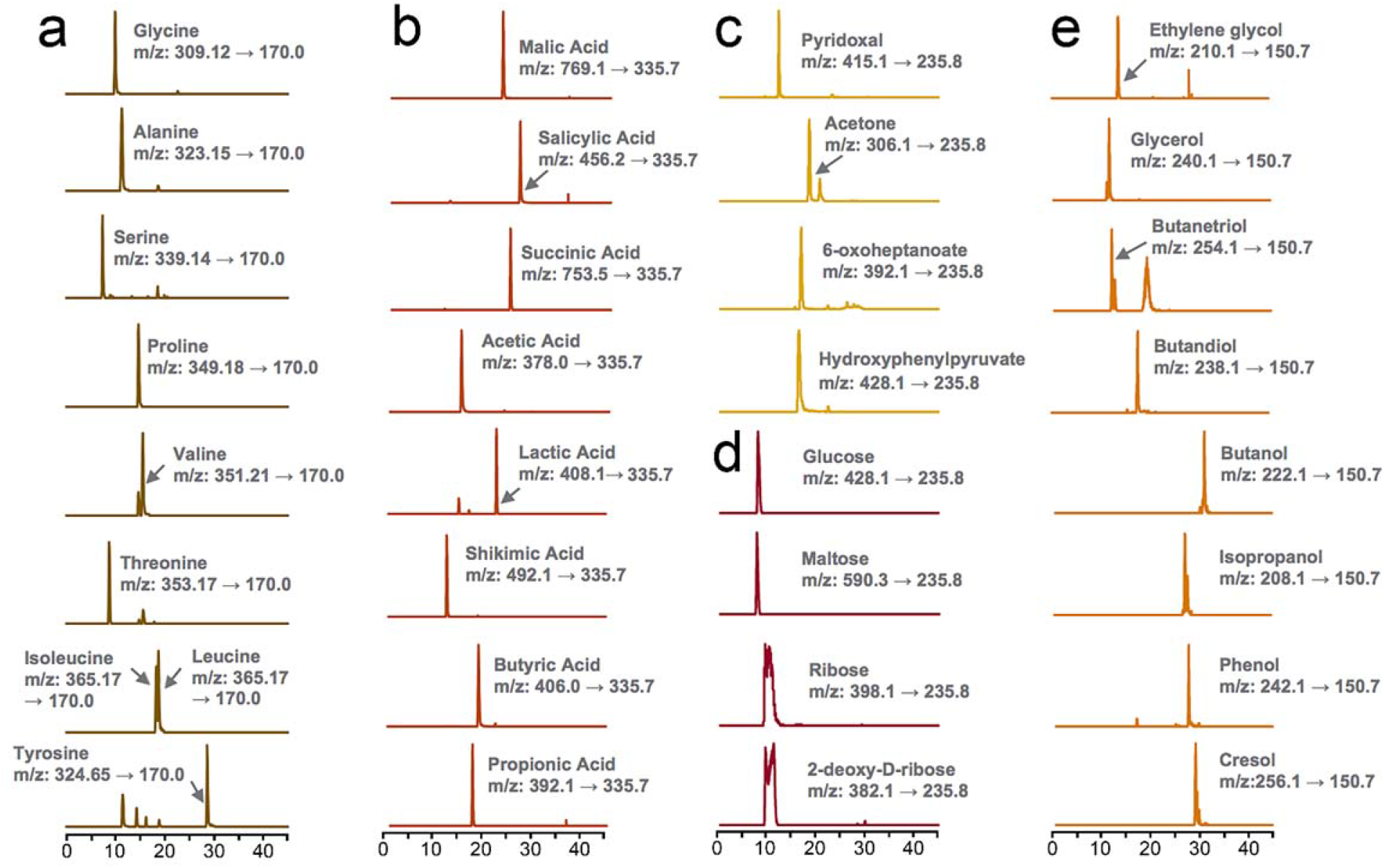
MetFish is applicable to measuring metabolites with a broad range of functional groups in challenging sample matrices. Shown are data from application of MetFish in measurement of (a) amine metabolites, (b) carboxyl metabolites, (c) carbonyl metabolites, (d) hydroxyl metabolites as sugars, and (e) hydroxyl metabolites as alcohols. In all cases, MetFish was deployed *in situ* in metabolite-salt mixtures containing 2 M MgSO4.

To further validate MetFish, we determined limits of quantification (LOQ), linear dynamic ranges, and relative standard deviations (RSD) for all four MetFish reagents and in measurements of 45 metabolites containing amine, carboxyl, carbonyl, or hydroxyl functional groups (**Supplemental Tables S3-6**) dissolved in water containing 2 M total salt. The amine-tagging method provided the lowest LOQ (median of 5 nM), the broadest linear dynamic range (5-6 orders of magnitude), and the lowest median inter-day reproducibility (median of 2.6%) of the four methods, based on data for 19 proteinogenic amino acids (**Supplemental Table S3**). The other tags showed median LOQs ranging from 40 nm (carboxyl; 10 metabolites) to 3.5 μM (hydroxyl; 8 metabolites), linear dynamic ranges of 3-5 orders of magnitude, and median interday RSD of 3.3% (carboxyl; 10 metabolites) to 9.3% (hydroxyl; 8 metabolites) (
**Supplemental Tables S4-6**). The hydroxyl tagging approach gave the highest LOQ, ranging from sub to low micromolar. All four MetFish tags showed excellent linearity over the dynamic range of quantification with R^2^ of 0.99 (**Supplemental Tables S3-6**).

### Application of MetFish in targeted analyses of proteinogenic amino acids in hypersaline matrices

After validating that the MetFish method can be used to enrich polar metabolites from a model hypersaline solution, we then applied the amine capture reagent in quantification of amino acids in exo-metabolomics analyses of two microbial communities: 1) a uni-cyanobacterial phototrophic microbial community and 2) a prairie soil.

MetFish was used to examine nitrogen metabolism over a 28-day succession in a unicyanobacterial consortial biofilm isolated from a benthic phototrophic microbial mat from a highly saline alkaline lake in northern Washington state.^38, 39,41^ During the seasonal cycle, the salt concentration in the lake fluctuates from low hundreds of mM to well over 2 M total dissolved salts (primarily MgSO_4_); the consortium in this experiment was therefore cultured in a defined medium containing 400 mM MgSO_4_ (see medium composition in **Supplemental Table S7** for full details). As organisms in the consortium are divergent for their ability to incorporate nitrate,^42^ this experiment aimed to determine how differences in the organismal access to nitrogen for amino acid biosynthesis influenced community dynamics and metabolite exchange. To test the hypothesis that availability of reduced nitrogen would increase the rate of amino acid sharing, the nitrate-containing growth medium was amended with either ammonium or urea. The samples were spiked with ^13^C and ^15^N-labeled amino acid standards, and endogenous amino acids in the media were quantified using isotope dilution MS. The MetFish analysis quantified 14 extracellular proteinogenic amino acids over a 17-day cultivation period (**Fig. 4a**). In general, amino acid concentrations increased to detectable levels early in cultivation until they reached a maximum at ~7-9 d, for nitrite and ammonium, or 4 d for urea, and decreased thereafter. Surprisingly, this trend did not hold true for all amino acids. For example, serine reached a maximum concentration at 14 d in medium amended with nitrate and at 11 d for ammonium. For proline, the maximum extracellular concentration occurred at 11 d for both ammonium and urea. The exo-metabolomics analysis of amino acid profiles during the phototrophic consortia succession revealed that availability of extracellular amino acids as community “public goods” differed among nitrogen sources at the level of individual amino acids. MetFish therefore enabled us to conclude that the nitrogen source for amino acid biosynthesis rewires overall community amino acid exchange.

**Figure 4.**
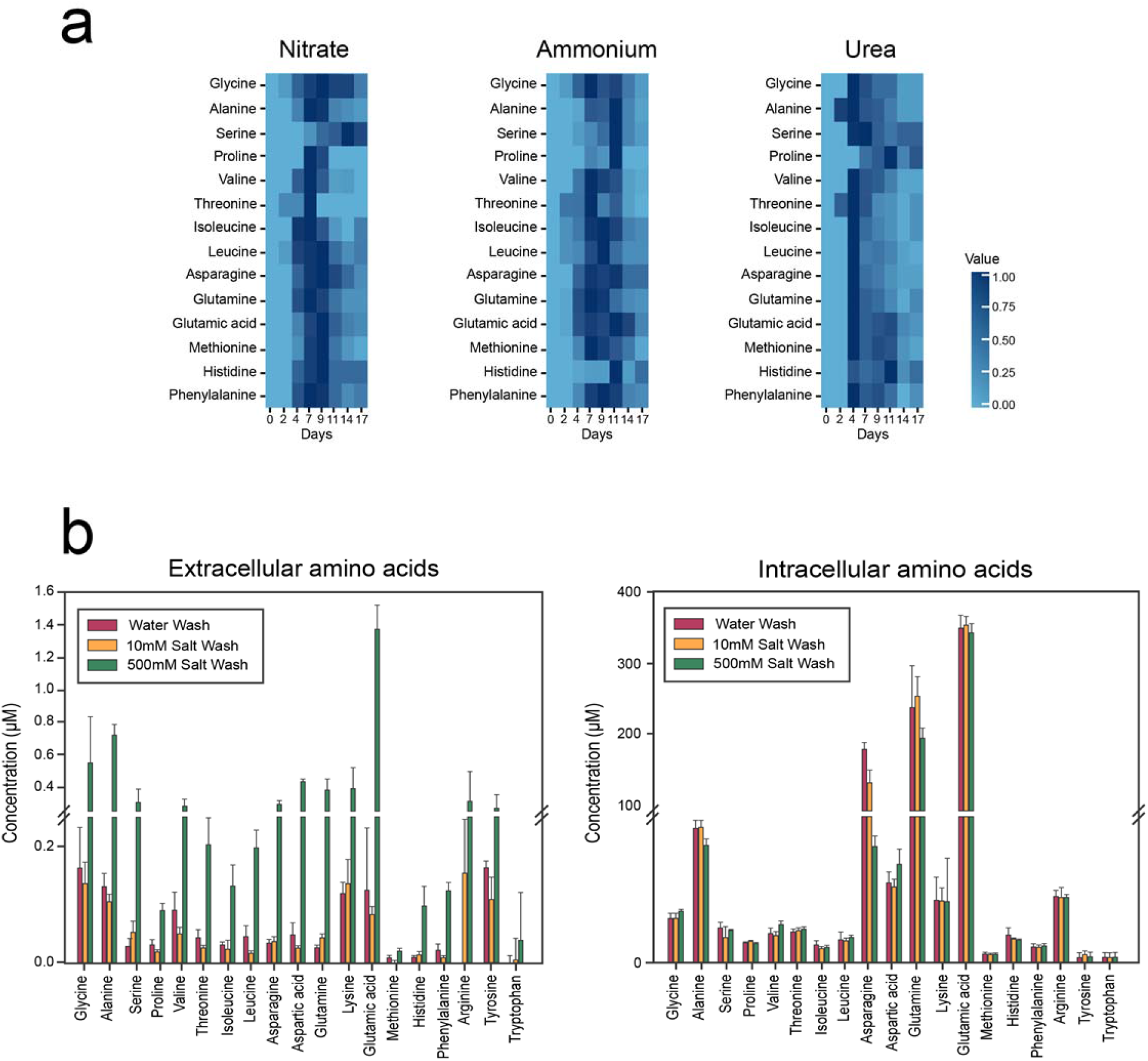
Application of MetFish in quantification of proteinogenic amino acids in representative microbial communities. (a) Quantification of amino acids during phototrophic microbial community succession with various nitrogen amendments (data shown are normalized average amino acid concentrations from analysis of 3 replicate succession experiments). (b) Exo-and endometabolomics analysis of amino acids in soil, using a high salt wash to increase recovery due to possible disruption of non-specific binding to soil particles (data shown are mean ± standard deviation from analysis of 3 replicate soil samples).

We next used MetFish in exo-metabolomics analyses to quantify free proteinogenic amino acids in soil, followed by analysis of biomass-associated molecules. To do so, we modified the classic fumigation-extraction method^43^ for measuring microbial biomass-associated carbon content. In the traditional format, soil samples are fumigated with chloroform to lyse microbial cells, followed by immediate extraction with 500 mM K_2_SO_4_, which extracts the total of free and biomass-associated molecules but cannot be used to distinguish between the two.^44^’^46^ Makarov and colleagues reported that microbial biomass-associated carbon is increasingly extractable with increasing concentration of the K_2_SO_4_ extraction solution, with solubility increases of 1.5 – 3.9-fold in 500 mM K_2_SO_4_ compared with 50 mM K_2_SO_4_.^47^ We therefore hypothesized that performing a 500 mM K_2_SO_4_ extraction of soil *prior* to microbial cell lysis would allow us to obtain higher recovery of molecules located in the extracellular milieu, and also enable us to follow up with a subsequent measurement of microbial biomass-associated molecules. Because the hypersaline environment of the salt extract would otherwise prohibit a LC-MS-based exo-metabolomics analysis, MetFish was employed. We used three different extractants – deionized water, 10 mM K_2_SO_4_, and 500 mM K_2_SO_4_ – to extract amino acids from a native prairie soil at the Konza Prairie Biological Station, a long-term ecological research site located in eastern Kansas, U.S.A. We extracted equivalent size aliquots of soil in replicate accordingly (see **Supplemental Methods** for details), and subsequently spiked the extracts with ^13^C and ^15^N-labeled amino acid standards and applied the amine-tagging MetFish reagent. The extracted soil remaining was then subjected to bead beating to lyse microbial cells, followed by spiking with labeled standards and derivatization of amino acids directly in the soil samples, demonstrating *in situ* applicability of MetFish. Nineteen proteinogenic amino acids in both the free and biomass-associated extracts were quantified using isotope dilution MS (**Fig. 4b**). As expected, pre-extraction of the soil with 500 mM K_2_SO_4_ resulted in 2-10-fold higher recovery of amino acids from the extracellular milieu compared to pre-extraction with water and 10 mM K2SO4. Asparagine, glutamine, glutamic acid were the three most abundant biomass-associated amino acids with concentrations of 70.9 μM/mg, 191.7 μM/mg, and 337.7 μM/mg soil, respectively (**Fig. 4b**). Intracellular levels of amino acids were similar between the three different pre-extractants.

### Application of MetFish in untargeted metabolomics analysis of fluids injected into and produced from a hydraulically fractured well

As described above, each MetFish reagent generates one or more tag-specific fragment ions during collision-induced dissociation during MS analysis. These “reporter ions” can be exploited in untargeted exo-metabolomics analyses to broadly query the metabolome in otherwise MS-prohibitive sample matrices. To demonstrate this, we applied all 4 MetFish reagents in parallel analyses of fluids injected into and produced from a hydraulically fractured well from the Utica-Point Pleasant shale (Ohio, U.S.A.), and operating the mass spectrometer in data-dependent MS/MS mode to obtain comprehensive untargeted data (**Fig 5a**). Although the complete composition of fracture fluid is typically proprietary, the fracking fluid used in our analyses was known to be complex, with up 125 g / L total dissolved solids, including salts, various corrosion inhibitors, and gelling agents. We initially applied all 4 MetFish reagents in untargeted exo-metabolomics analysis of a representative produced fluid sample, in order to identify as many putative molecules as possible (see **Supplemental Methods** for details) (**Fig 5b**). We then purchased isotopically-labeled standards for putatively identified metabolites, and applied MetFish in a targeted exo-metabolomics analysis to confirm molecular identities in a time series of produced fluid samples ranging from 86 and 154 d post-injection (**Fig. 6a**). Using this approach, we confirmed the identities of 37 metabolites. As shown in **Fig. 6a**, fluids initially produced from the well at 86-98 d showed higher amounts of amino acids than those at 105 and 154 d, while the concentrations of most alcohols and organic acids detected were evenly distributed over the time course. Compared to the input fluids, the metabolite concentrations in produced fluid samples show significant differences (**Fig. 6b**). Metabolites such as amino acids and organic acids have significantly higher concentrations in produced fluid samples than in the input fluids, indicating the presence of metabolically active microbial communities. The input fluids also contained extremely high levels of diols, such as propylene glycol, which are typical additives in hydraulic fracture fluids.

**Figure 5.**
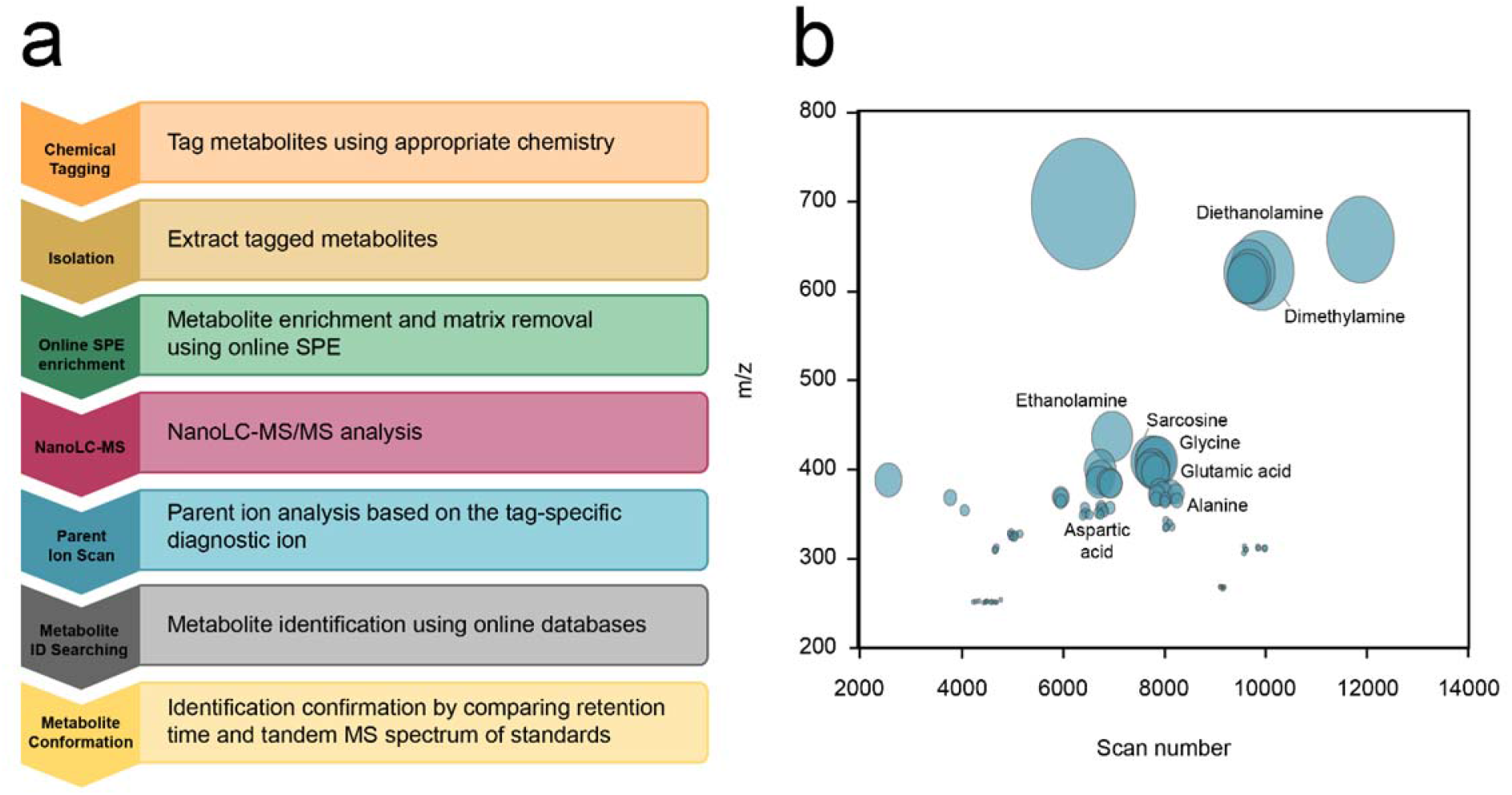
(a) Workflow for untargeted metabolomics analysis using MetFish. (b) Global metabolite profile of a produced fluid sample. Data from all 4 MetFish reagents were combined into a single plot of *m/z* vs MS scan number. The size of the circle is proportional to the ion intensity, and putatively identified metabolites are labeled.

**Figure 6.**
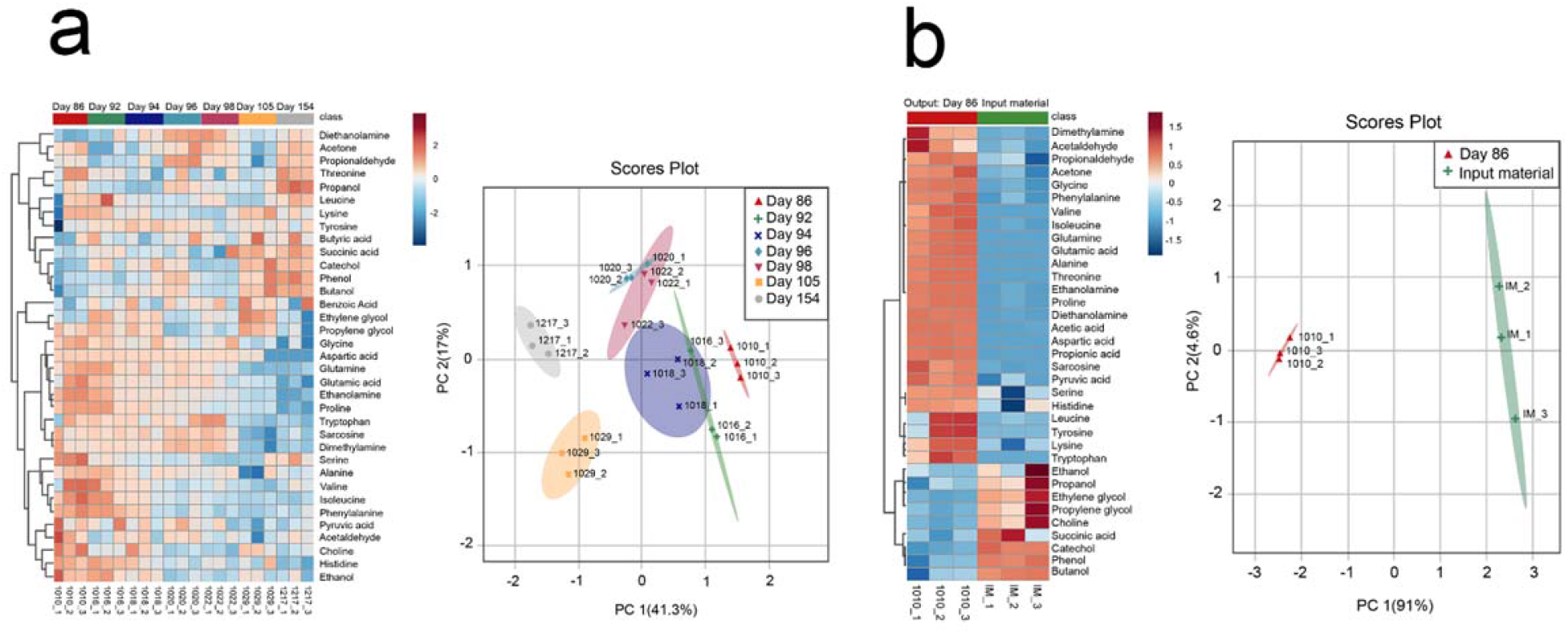
Targeted metabolomics analysis using MetFish in injected and produced fluids from hydraulic fracturing. **(a)** Quantification of 37 metabolites identified by targeted MetFish **(b)** Comparison of metabolite levels in the input material and spent fracking fluid. Data shown are from replicate analysis (n = 3) of each fluid sample.

## Discussion

In summary, the MetFish method enables highly sensitive targeted and untargeted exo-metabolomics measurements in chemically extreme environments that are otherwise prohibitive to MS-based analyses. We demonstrated use of MetFish in quantification of exo-metabolites in hypersaline matrices, including spent media from a phototrophic microbial consortium, salt-extracted soil, and injected/produced fluids from hydraulic fracturing. The combination of a high salt wash and MetFish was particularly useful for extracting metabolites from the extracellular soil milieu, prior to subsequent *in situ* application of MetFish for analysis of intracellular metabolites in the same samples after microbial cell lysis. The use of MetFish offers control over the sub-class of metabolites being captured, which greatly constrains the chemical search space when attempting to identify unknowns during untargeted exo-metabolomics analysis. This is particularly useful for samples containing a diversity of high concentration organic constituents, such as soils or those produced from hydrocarbon-bearing, hydraulically-fractured wells. We believe that such an approach will aid in the investigation of metabolite exchange in microbial communities and provide a more effective way to understand the microbial metabolism in extreme ecosystems that remain understudied.

## Supporting information

Supplemental Section

## Acknowledgements

This research was supported by the Genomic Science Program (GSP), Office of Biological and Environmental Research (OBER), U.S. Department of Energy (DOE), and is a contribution of the Metabolic and Spatial Interactions in Communities Scientific Focus Area. Additional support was provided by the Pacific Northwest National Laboratory (PNNL), Laboratory Directed Research and Development program via the Microbiomes in Transition Initiative. P.J.M was partially supported by funding from the National Sciences Foundation Dimensions of Biodiversity (award no. 1342701). The authors thank Susan Welch with the Ohio State University Subsurface Energy Materials Characterization and Analysis Laboratory (SEMCAL) for field sampling assistance as well as Kelly Wrighton, Mike Wilkins, and Kayla Borton for providing samples and their preparation; Dr. Aaron Wright of PNNL for useful discussions and assistance with the MetFish derivatization reactions; and Drs. Charles Rice and Ari Jumpponen at Kansas State University for providing the soil samples. Targeted and untargeted metabolomics analyses using MetFish were performed in the Environmental Molecular Sciences Laboratory, a national scientific user facility sponsored by the U.S. OBER and located at PNNL in Richland, Washington. PNNL is a multi-program national laboratory operated by Battelle for the DOE under Contract DE-AC05-76RL0 1830.

## Author contributions

C.X. and T.O.M. conceived and designed the method and studies. S.R.L., T.R.C., R.Z., R.J.M., J.K.J., V.L.B., P.J.M., and M.F.R. contributed materials and assisted with experimental design. C.X., R.L.S., N.G.I., Y.M., and B.R.M. performed experiments and data analysis. M.F.R. and J.K.F. provided funding and critical review of the manuscript. C.X., S.P.C. and T.O.M. performed data analysis, interpreted results, and wrote the manuscript. All authors read and approved the final manuscript.

## Data availability

The data that support the findings of this study are available upon request.

