## Supplemental Section for "MetFish: A Metabolomics Platform for Studying Microbial Communities in Chemically Extreme Environments"

### Supplemental Information

#### Methods

##### MetFish chemical tagging methods

**Amine reagent**. 200 µL of amine-containing metabolites in water, media or matrix were combined with 500 µL of dansyl chloride in acetonitrile (40 mM) and diluted to a final volume of 1200 µL with 0.5 M $Na_2CO_3$-$NaHCO_3$ buffer (pH 9.5). The reaction solution was mixed using a Thermomixer (Eppendorf, Hauppauge, NY, USA) at 60°C for 40 min at 1500 rpm. The organic portion of the resulting solution was removed using a SpeedVac (Eppendorf) for 30 min, and the pH of the aqueous portion was adjusted to 3-4 by using 20% formic acid (v/v). Then a liquid-liquid extraction was performed using dichloromethane and water. The organic layer was collected and reconstituted in 1 mL of a mixture of water and methanol (95:5, v/v) containing 0.1 % of formic acid followed by online SPE-nanocapillary LC-MS/MS analysis.

**Carbonyl reagent**. 200 µL of carbonyl-containing metabolites in water, media or matrix were combined with 50 µL of 10% trichloroacetic acid in water (w/v), 250 µL dansyl hydrazine in ethanol (10 mM), and diluted to a final volume of 900 µL with 400 µL of water. The reaction solution was mixed using a Thermomixer (Eppendorf, Hauppauge, NY, USA) at 60°C for 15 min at 1500 rpm. The organic portion of the resulting solution was removed using a SpeedVac (Eppendorf) for 30 min, and the pH of the aqueous portion was adjusted to 3-4 by using 20% formic acid (v/v). A liquid-liquid extraction was then performed by using dichloromethane and water. The organic layer was collected and reconstituted in 1 mL of a mixture of water and methanol (95:5, v/v) containing 0.1 % of formic acid followed by online SPE-nanocapillary LC-MS/MS analysis.

**Carboxylic acid reagent**. 150 µL of carboxylic acid-containing metabolites in water, media, or matrix were combined with 300 µL of 1-ethyl-3-(3-dimethylaminopropyl)carbodiimide in water

(10 mM), 300 µL dansylcadaverine in methanol (10 mM), 300 µL hydroxybenzotriazole in ethanol, and diluted to a final volume of 1350 µL with 300 µL of water. The reaction solution was mixed using a Thermomixer (Eppendorf, Hauppauge, NY, USA) at 60°C for 120 min at 1500 rpm. The organic portion of the resulting solution was removed using a SpeedVac (Eppendorf) for 30 min, and the pH of the aqueous portion was adjusted to 3-4 by using 20% formic acid (v/v). Then a liquid-liquid extraction was performed using dichloromethane and water. The organic layer was collected and reconstituted in 1 mL of a mixture of water and methanol (95:5, v/v) containing 0.1 % of formic acid followed by online SPE-nanocapillary LC-MS/MS analysis.

**Hydroxyl reagent**. 300 µL of hydroxyl-containing metabolites in water, media, or matrix were combined with 300 µL 4-dimethylaminobenzoic chloride in tetrahydrofuran (10 mM), and diluted to a final volume of 900 µL with 300 µL of 100 mM sodium carbonate in water. The reaction solution was mixed using a Thermomixer (Eppendorf, Hauppauge, NY, USA) at 30°C for 50 min at 1500 rpm. The organic portion of the resulting solution was removed using a SpeedVac (Eppendorf) for 30 min, and the pH of the aqueous portion was adjusted to 3-4 using 20% formic acid (v/v). Then a liquid-liquid extraction was performed using dichloromethane and water. The organic layer was collected and reconstituted in 1 mL of a mixture of water and methanol (95:5, v/v) containing 0.1 % of formic acid followed by online SPE-nanocapillary LC-MS/MS analysis.

#### Sample preparation for metabolomics analyses

##### Photoautotrophic microbial consortium

The photoautotrophic microbial consortium was routinely cultured, as described previously. Briefly, biofilms were grown and maintained in T75 Corning™ Cell Culture Flasks (Fisher # 10-126-37) on Hot Lake Autotroph medium (HLA; see Table 7S for details) at room temperature and atmosphere under ~35 $\mu E/m^2/s$ light (PL/AQ; General Electric) over 28-day succession experiments. Sterile water was replaced weekly to replace volume lost to evaporation. For exometabolomics analysis of, 20 mL of cell culture was centrifuged at 6000 × g for 15 min at 4 °C. The supernatant was then transferred into a separated vessel. A portion of the supernatant was spiked with isotopically labeled amino acids as internal standards and analyzed using the amine tagging protocol.

**Fracture Fluid and Produced Fluids**

Due to the high acidity of fracking fluid samples, the samples were pretreated by adding 250 µL of 0.5 M NaOH solution. Then, the MetFish methodology was applied to the fracture fluid and produced fluid samples by using each chemical tagging approach followed by LC-MS/MS analysis. A Q Exactive Hybrid Quadrupole-Orbitrap high resolution mass spectrometer (Thermo Fisher Scientific, San Jose, CA) was used for metabolite profiling, and data-dependent MS/MS spectra were obtained. Data analysis was performed by using an in-house developed software MASIC to extract tag-derived metabolite masses with the use of three diagnostic fragments for each tagging approach. The masses of unknown metabolites were matched against the METLIN database and the Human Metabolome Database (HMDB) using a mass accuracy < 10 ppm.

**Soil**

**Extracellular soil metabolite extraction.** Silty-loam soil was collected from the upper 15-cm of a watershed (39˚06’11" N, 96˚36’48" W) located at the Konza Prairie Biological station. Upon collection, soil samples were shipped on ice to the Pacific Northwest Laboratory, where soils were immediately sieved (< 2mm) and stored at -80 $^{o}$C until further use. Soil pH (in water) was 6.5 and sulphate concentration was 5.45 ppm. Soil gravimetric water and clay content were determined to be 37 and 2%, respectively. A salt wash of soil samples was performed in order to completely extract extracellular metabolites from soil particulates. 500 mM and 10 mM $K_2SO_4$ solutions were used for salt washes; a water extraction was also performed as control. Each extraction was performed in triplicate. 1 g of sieved soil sample and 2 mL $K_2SO_4$ salt wash solution or water were added into a centrifuge tube. The mixture was vortexed for 30 min. The resulting mixture was centrifuged at 5000 rpm for 5 min and the supernatant was depleted. Then another salt or water wash was performed on the residual soil sample. Another 2 mL $K_2SO_4$ salt wash solution or water was added to the residual soil sample and the mixture was vortexed for 30 min. The two supernatants were combined and was filtered through a 0.22 µM filter. 20 µL of 300 µg/mL $^{13}$C, $^{15}$N-labeled amino acid mixture standard was added to the filtered supernatant. Then, the amine-specific tagging approach was conducted to analyze extracellular metabolites.

**Intracellular soil metabolite extraction.** The residual soil from the extracellular metabolite extraction was used for subsequent intracellular metabolite extraction. A scoop of stainless steel beads and garnet beads were added into the centrifuge tube. 1 mL of water was then added and bead beating was performed at speed 7 for 4 min at 4 °C to lyse microbial cells. 20 µL of 300 µg/mL $^{13}C$, $^{15}N$-labeled amine acid mixture standard was added to the sample. Then the amine-specific tagging approach was directly conducted in soil to quantify released intracellular amino acids.

#### Instrumentation

**Online SPE-nanocapillary liquid chromatography system**

An online SPE-nanocapillary liquid chromatography system was used for analysis of MetFish tagged metabolites. The system consists of two parallel subsystems, each of which consists of two Agilent 1200 series nano pumps (Agilent Technologies, Santa Clara, CA), a six-port injection valve (VICI Valco, Houston, TX) with a sample loop, and a six-port injection valve (VICI Valco, Houston, TX) with a micro solid phase extraction (SPE) column coupled in line **(Supplemental Figure S1)**. The valves were switched by program and directed the LC flow to carry the analyte from the sample loop into the SPE. After the analyte was enriched at SPE, it was backflushed to a nine-port valve (VICI Valco, Houston, TX), which delivered the analyte into the nanocapillary LC column. The nanocapillary LC column was packed with porous C18 particles (3 µm particle size, 300 Å pore size, Phenomenex, Terrence, CA) in fused silica capillaries (35 cm × 75 µm i.d. × 360 µm o.d., Polymicro Technologies, Phoenix, AZ). The outlet of the nanocapillary LC column was connected to an approximately 3-cm-long nanoESI emitter, which was chemically etched from a 20 µm i.d. × 150 µm o.d. fused silica capillary. An autosampler (LEAP Technologies, Carrboro, NC) with a cooled-drawer sample holder was used for both parallel subsystems. A 5 µL sample loop was used for all experiments except noted otherwise. Chromatography solvents consisted of 0.1 % formic acid in water (mobile phase A) and 0.1 % formic acid in acetonitrile (mobile phase B). Chromatographic separation was performed by using gradient elution of 2-30% B in 5 min, 30- 95% B in 35 min, and 95% B for

20 min. The LC system was operated at constant flowrate at 300 nL/min. Data acquisition begins after 10 min of gradient elution to avoid recording of data-poor regions.

**Mass spectrometry**

A TSQ Quantum Ultra mass spectrometer (Thermo Fisher Scientific, San Jose, CA) was used for targeted metabolite quantification. The mass spectrometer was operated with an electrospray voltage of +2400 V, a capillary offset voltage of 35V, an ion transfer capillary temperature of 310 °C, and a skimmer offset voltage of 0 V. Tube lens voltages were obtained from automatic tuning without further optimization. In selected reaction monitoring (SRM) mode, both Q1 and Q3 were set at a resolution of 0.7 FWHM and Q2 gas pressure was set at 1.5 mTorr. Scan width was set at 0.002 m/z and dwell time was set at 25 ms. Scan time was typically set at 20 ms for each SRM transition and adjusted as necessary based on transition numbers to provide enough data points across the chromatographic peaks. The collision energy was optimized for each SRM transition by using automatic tuning. Peak identification and quantification was performed by using Quan Browser provided by the Xcalibur 2.0 software.

A Q Exactive Hybrid Quadrupole-Orbitrap high resolution mass spectrometer (Thermo Fisher Scientific, San Jose, CA) was used for untargeted metabolite analysis. The mass spectrometer conditions were as follows: electrospray voltage, +3900 V; ion transfer capillary temperature, 320 °C; source DC offset, 21V; and S-lens RF level, 50. The instrument was operated in data-dependent mode acquiring high-resolution full scan (Resolution = 70 000, AGC = $3 \times 10^6$) spectra followed by MS/MS scans (Resolution = 70 000, AGC = $1 \times 10^5$) of the top five most abundant ions within the mass range of 200 to 2000 m/z. An isolation window of 2.0 Da was used. The dynamic exclusion function was not used. Stepped normalized collision energy (NCE) during collision induced dissociation (CID) was set as 30, 33, 36. Data were process by using an in-house tool MASIC. The reporter ions for each chemically tagging approach were identified and the corresponding parent ion mass spectra were generated by using MASIC.

**Analytical method validation**

Fragment ions for each analyte were generated from product ion scans using the TSQ Quantum Ultra mass spectrometer (Thermo Fisher Scientific, San Jose, CA). The most abundant ion generated from during collision-induced fragmentation was selected as the quantification ion, and another two abundant and characteristic fragment ions were used as confirmation ions. The quantification of metabolites was accomplished in the SRM mode by using internal calibration ($R^2$>0.99). The calibration curve was produced by plotting the ratio of peak area of analyte to peak area of the internal standard versus the concentration of metabolite standard solution before derivatization. The method detection limits were determined by dilution series of metabolite standard solutions followed by chemical tagging until a signal-to-noise ratio of 10 was noted in LC-MS chromatogram. The signal-to-noise ratio was calculated based on a Genesis Peak Detection Algorithm provided by the Xcalibur 2.0 software. Reproducibility was examined by evaluating the relative standard deviation (RSD) for three days (inter-day reproducibility) using sample solutions spiked with 5 µM of metabolite standards.

#### Testing the efficacy of SPE columns for enriching metabolites from hypersaline media

To evaluate the efficacy of SPE columns for enriching metabolites from hypersaline media, metabolites (see **Supplemental Table S1**) were combined at 500 µM in both water and 0.5 M $K_2SO_4$. SPE columns were conditioned, loaded, washed, and metabolites extracted as listed below. Protocols were adapted from multiple sources, including manufacturer recommendations, published literature, and previous experience. Bed volumes for C-18, SXC, and SAX columns were 50 mg, while graphitic carbon was 100 mg. After extraction, samples were dried to completeness in a Speedvac and re-suspended in 1 ml of water. 270 µl of resuspended sample was combined with 30 µl of DSS (NMR reference standard) and placed in a 3 mm NMR tube for analysis.

Data were acquired on a Varian Direct Drive (VNMRS) 600 MHz spectrometer (Agilent Technologies). The spectrometer system was outfitted with a Varian triple resonance salt-tolerant cold probe with a cold carbon preamplifier. A Varian standard one dimensional proton nuclear Overhauser effect spectroscopy (noesy) with presaturation was collected on each sample, using a nonselective 90 degree excitation pulse, a 100 millisecond mixing time, acquisition time of 4 seconds, a presaturation delay of 1.5 seconds, spectral width of 12 ppm, and temperature control set to 25C. Collected spectra were analyzed using Chenomx 8.3 software, with quantifications based on spectral intensities relative to 0.5 mM 3-(trimethylsilyl)-1-propanesulfonic acid-d6 (Sigma-Aldrich), which was added as a spike to each sample.

### Supplemental Figures and Tables

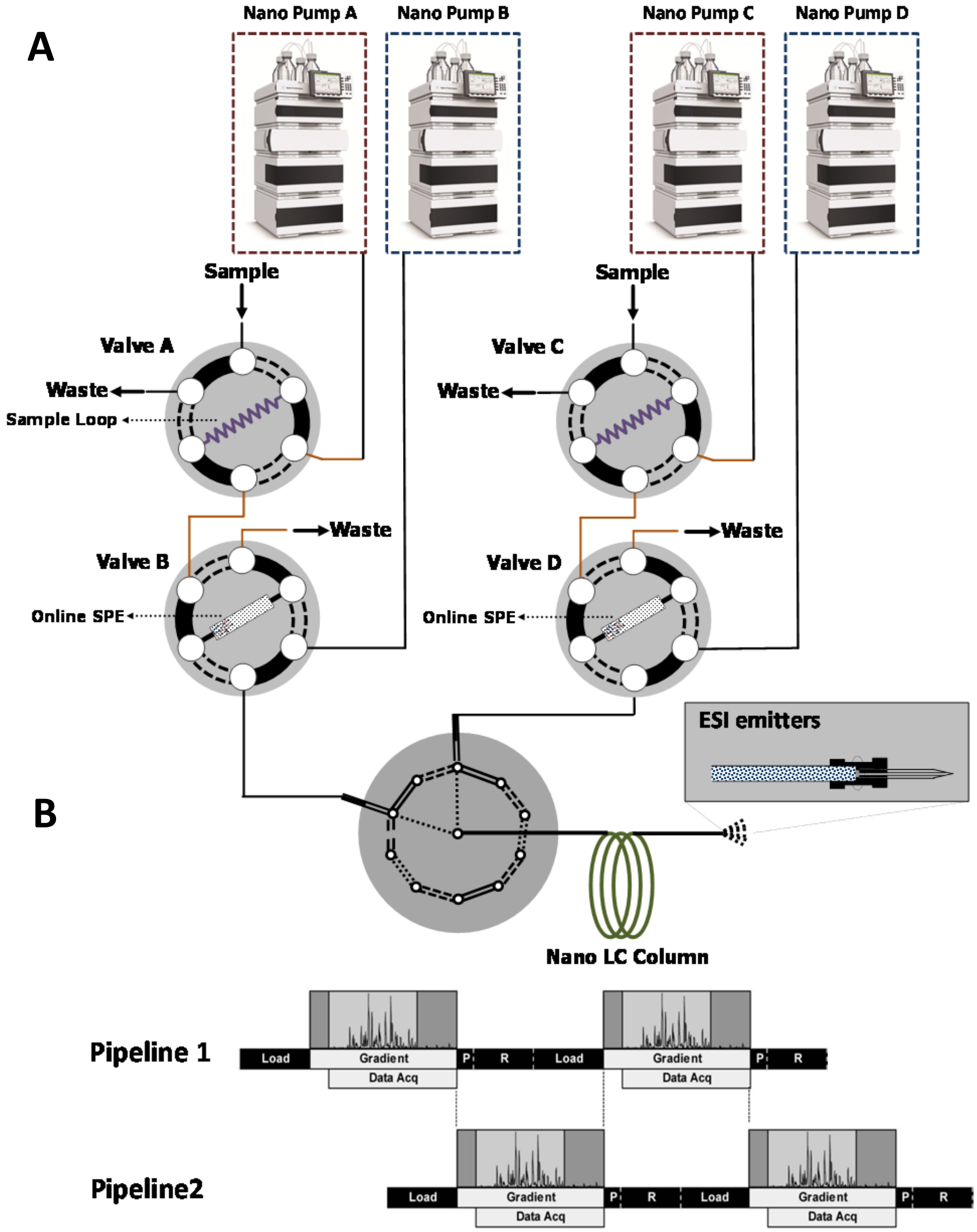


**Figure S1**. (**A**) Instrumental setup of MetFish methodology. (**B**) Operation timeline of the two parallel pipelines.

**Table S1.** Performance of commercial SPE columns for separation of metabolites from high salt matrices, as determined by NMR spectroscopy. Generally, SPE columns were not effective in recovering metabolites from either water or hypersaline media, although some column chemistries were effective to some extent for certain metabolites.

| Name | Starting Amount | | % Recovery in eluent | | | | | | | |
|---|---|---|---|---|---|---|---|---|---|---|
| | $H_2O$ | $K_2SO_4$ | $H_2O$ (SAX) | $K_2SO_4$ (SAX) | $H_2O$ (SCX) | $K_2SO_4$ (SCX) | $H_2O$ (Carbon) | $K_2SO_4$ (Carbon) | $H_2O$ (C-18) | $K_2SO_4$ (C-18) |
| 2-Oxoglutarate | 262.4 | 393.3 | 0.00 | 0.00 | 0.00 | 0.00 | 16.04 | 0.00 | 0.00 | 0.00 |
| Acetate | 12.6 | 21.3 | 451.59 | 532.39 | 34.92 | 21.13 | 1137.30 | 470.42 | 61.60 | 30.60 |
| Alanine | 444.4 | 488.1 | 0.00 | 0.00 | 0.00 | 0.00 | 0.00 | 0.00 | 0.00 | 0.00 |
| Aspartate | 454.8 | 535.6 | 0.00 | 0.00 | 0.00 | 0.00 | 0.00 | 0.00 | 0.00 | 0.00 |
| Fumarate | 703.4 | 781.4 | 0.00 | 0.00 | 0.00 | 0.00 | 13.08 | 1.51 | 0.00 | 0.00 |
| Glucose | 501.1 | 581.4 | 0.00 | 0.00 | 0.00 | 0.00 | 0.00 | 0.00 | 0.00 | 0.00 |
| Lysine | 329.2 | 421.3 | 0.00 | 0.00 | 0.00 | 0.00 | 0.00 | 0.00 | 0.00 | 0.00 |
| Phenylalanine | 728.1 | 755 | 0.00 | 0.00 | 0.00 | 0.00 | 90.77 | 89.06 | 0.00 | 7.10 |
| Phosphoenolpyruvate | 2027 | 2702.4 | 2.49 | 0.00 | 0.00 | 0.00 | 0.00 | 0.00 | 0.00 | 0.00 |
| Proline | 500.7 | 488.4 | 0.00 | 0.00 | 0.00 | 0.00 | 0.00 | 0.00 | 0.00 | 0.00 |
| Pyruvate | 40.8 | 114.8 | 0.00 | 0.00 | 0.00 | 0.00 | 12.99 | 0.00 | 0.00 | 0.00 |
| Succinate | 530.1 | 586.9 | 0.00 | 0.00 | 0.00 | 0.00 | 5.38 | 0.00 | 0.00 | 0.00 |
| Trehalose | 530 | 598.1 | 0.00 | 2.99 | 0.00 | 0.00 | 35.74 | 32.79 | 0.00 | 0.00 |
| Uracil | 688.6 | 760.1 | 0.00 | 0.00 | 0.00 | 0.00 | 71.07 | 66.21 | 0.00 | 0.00 |
| Valine | 445.1 | 475.4 | 0.00 | 0.00 | 0.00 | 0.00 | 0.00 | 0.00 | 0.00 | 0.00 |

**Table S2.** Common fragment ions generated during MS/MS for MetFish reagents.

| MetFish reagent | Common fragment ions | | |
|---|---|---|---|
| | **Fragment Ion 1** | **Fragment Ion 2** | **Fragment Ion 3** |
| Amine tagging reagent | 170.1 | 156.6 | 234.1 |
| Carboxylic tagging reagent | 170.1 | 335.8 | 234.1 |
| Carbonyl tagging reagent | 157.1 | 170.1 | 235.8 |
| Hydroxyl tagging reagent | 133.6 | 151.1 | 170.1 |

**Table S3.** Limits of quantification (LOQ), linearity, and interday reproducibility of amine tagging approach.

| Metabolite | LOQ (nM) | Linear Dynamic Range | $R^2$ | Interday Reproducibility (RSD%) |
|---|---|---|---|---|
| Glycine | 4 | 4 nM - 1 mM | 0.99 | 5.0 |
| Alanine | 9 | 9 nM – 2.5 mM | 0.99 | 2.4 |
| Serine | 30 | 30 nM - 10 mM | 0.99 | 1.5 |
| Proline | 4 | 4 nM – 2.5 mM | 0.99 | 1.1 |
| Valine | 5 | 5 nM - 1 mM | 0.99 | 1.3 |
| Threonine | 20 | 20 nM – 2.5 mM | 0.99 | 1.9 |
| Isoleucine | 3 | 4 nM – 0.2 mM[a] | 0.99 | 3.9 |
| Leucine | 2 | 4 nM – 0.2 mM[a] | 0.99 | 4.8 |
| Asparagine | 20 | 20 nM – 2.5 mM | 0.99 | 2.6 |
| Aspartic acid | 40 | 40 nM – 2.5 mM | 0.99 | 2.6 |
| Glutamine | 20 | 20 nM – 2.5 mM | 0.99 | 1.6 |
| Lysine | 4 | 4 nM – 2.5 mM | 0.99 | 6.4 |
| Glutamic acid | 40 | 40 nM – 2.5 mM | 0.99 | 1.6 |
| Methionine | 1 | 1 nM – 0.5 mM | 0.99 | 2.1 |
| Histidine | 4 | 4 nM – 0.5 mM | 0.99 | 5.0 |
| Phenylalanine | 2 | 2 nM – 2.5 mM | 0.99 | 4.6 |
| Arginine | 90 | 90 nM – 2.5 mM | 0.99 | 6.8 |
| Tyrosine | 80 | 80 nM – 2.5 mM | 0.99 | 8.7 |
| Tryptophan | 3 | 3 nM – 2.5 mM | 0.99 | 2.5 |

[a] when the concentrations of isoleucine and leucine exceed 20 µM, they cannot be resolved ($R_s < 1.0$).

**Table S4.** Limits of quantification (LOQ), linearity, and interday reproducibility of carboxylic acid tagging approach.

| Metabolite | LOQ (nM) | Linear Dynamic Range | $R^2$ | Interday Reproducibility (RSD%) |
|---|---|---|---|---|
| Butyric Acid | 20 | 20 nM – 2500 µM | 0.99 | 0.4 |
| Propionic Acid | 80 | 80 nM – 1000 µM | 0.99 | 2.0 |
| Acetic Acid | 180 | 180 nM – 1000 µM | 0.99 | 2.2 |
| Salicylic Acid | 30 | 30 nM – 2500 µM | 0.99 | 4.4 |
| Shikimic acid | 20 | 20 nM – 2500 µM | 0.99 | 13.9 |
| Lactic Acid | 70 | 70 nM – 1000 µM | 0.99 | 1.7 |
| Pyruvic Acid | 30 | 30 nM – 100 µM | 0.99 | 1.3 |
| Oxalic Acid | 50 | 50 nM – 500 µM | 0.99 | 5.7 |
| Malic Acid | 10 | 10 nM – 500 µM | 0.99 | 9.6 |
| Succinic Acid | 100 | 100 nM – 100 µM | 0.99 | 4.4 |

**Table S5.** Limits of quantification (LOQ), linearity, and interday reproducibility of carbonyl tagging approach.

| Metabolite | LOQ (nM) | Linear Dynamic Range | $R^2$ | Interday Reproducibility (RSD%) |
|---|---|---|---|---|
| 6-oxoheptanoate | 300 | 300 nM – 2000 µM | 0.99 | 4.4 |
| Hydroxyphenyl pyruvate | 300 | 300 nM – 500 µM | 0.99 | 9.6 |
| Pyridoxal | 2000 | 2000 nM – 1000 µM | 0.99 | 14.7 |
| Acetone | 5000 | 5000 nM – 10000 µM | 0.99 | 12.5 |
| Glucose | 70 | 70 nM – 1000 µM | 0.99 | 5.3 |
| Maltose | 100 | 100 nM – 1000 µM | 0.99 | 12.3 |
| Ribose | 300 | 300 nM – 500 µM | 0.99 | 7.2 |
| 2-deoxy-D-ribose | 300 | 300 nM – 5000 µM | 0.99 | 6.5 |

**Table S6.** Limits of quantification (LOQ), linearity, and interday reproducibility of hydroxyl tagging approach.

| Analyte | LOQ (nM) | Linear Dynamic Range | $R^2$ | Interday Reproducibility (RSD%) |
|---|---|---|---|---|
| ethylene glycol | 3000 | 3000 nM – 5000 µM | 0.99 | 9.1 |
| glycerol | 2000 | 2000 nM – 5000 µM | 0.99 | 7.2 |
| 1,2,4-butanetriol | 700 | 700 nM – 5000 µM | 0.99 | 5.6 |
| 2,3-butandiol | 700 | 700 nM – 5000 µM | 0.99 | 9.6 |
| butanol | 5000 | 5000 nM – 5000 µM | 0.99 | 14.4 |
| isopropanol | 4000 | 4000 nM – 5000 µM | 0.99 | 10.7 |
| Phenol | 5000 | 5000 nM – 2000 µM | 0.99 | 7.8 |
| m-cresol | 5000 | 5000 nM – 2000 µM | 0.99 | 13.0 |

**Table S7. HLA medium composition for culturing of photoautotrophic microbial community.**

| Compound | CAS No. | Molecular Weight | Concentration |
|---|---|---|---|
| $MgSO_4 \cdot 7H_2O$ | 10034-99-8 | 246.48 g/mol | 400 mM |
| $Na_2SO_4$ | 7757-82-6 | 142.04 g/mol | 80 mM |
| KCl | 7447-40-7 | 74.55 g/mol | 20 mM |
| $NaNO_3$ | 7631-99-4 | 85 g/mol | 17.6 mM |
| TES Na salt (pH 8.0) | 70331-82-7 | 251.2 g/mol | 10 mM |
| $NaHCO_3$ | 144-55-8 | 84.01 g/mol | 1 mM |
| $CaCl_2 \cdot 2H_2O$ | 10035-04-8 | 147.01 g/mol | 245 µM |
| $Na_2CO_3 \cdot H_2O$ | 11/6/5968 | 124 g/mol | 189 µM |
| $K_2HPO_4$ | 11/4/7758 | 174.2 g/mol | 175 µM |
| $H_3BO_3$ | 10043-35-3 | 61.83 g/mol | 46.3 µM |
| $NH_4$ Fe(III) citrate | 1185-57-5 | 265 g/mol | 23 µM |
| $MnCl_2 \cdot 4H_2O$ | 13446-34-9 | 198 g/mol | 9.14 µM |
| $Na_2$EDTA | 139-33-3 | 372.24 g/mol | 2.8 µM |
| $Na_2MoO_4 \cdot 2H_2O$ | 10102-40-6 | 241.95 g/mol | 1.61 µM |
| $ZnSO_4 \cdot 7H_2O$ | 7446-20-0 | 287.55 g/mol | 720 nM |
| $CuSO_4 \cdot 5H_2O$ | 7758-99-8 | 249.68 g/mol | 316 nM |
| $Co(NO_3)_2 \cdot 6H_2O$ | 10026-22-9 | 291.03 g/mol | 169.7 nM |
